# Evidence of the Cost-Efficiency of Scale as seen in Polio Vaccination and Surveillance Costs

**DOI:** 10.1101/654350

**Authors:** Brittany L. Hagedorn, Laina D. Mercer, Guillaume Chabot-Couture

## Abstract

This analysis examined how polio program costs vary with scale for vaccination and disease surveillance, based on historical budget data published by the Global Polio Eradication Initiative (GPEI) from 2005 to 2018. We applied a linear mixed effects regression model in order to understand the cost structure of the historical GPEI budgets, with the goal that lessons learned from polio may be extended to other global disease elimination programs. Our findings demonstrate that there are economies of scale for vaccine delivery operations and for disease surveillance, which means that larger programs can leverage fixed costs and achieve better cost-efficiency as they scale. This finding should enable decision makers to create more reliable budgets, which support fundraising and optimal resource allocation. They also provide insight into how cost effectiveness changes as programs scale up during progressive disease control and elimination, as well as what level of resources are needed to sustain a program that is scaling back post-eradication and through to certification.

## Introduction

In earlier studies (1–3), vaccine delivery costs are reported on an average per-unit basis, such as per-vaccine or per-child. Averages are calculated as the total of many costs including categories such as salaries, transportation, vaccine acquisition, and supply chain, divided by the number of units (e.g. population targeted or the number of vaccines administered), such as in Walker et al. (2004) and Griffiths et al. (2016). Some analyses (6) include a sensitivity analysis and demonstrate that the average cost may vary from the reported value, but this only emphasizes the uncertainty in the point estimate, not that the drivers of cost may change structurally under different conditions.

In addition, Geng et al. (2017) and Schütte et al. (2015) have demonstrated that program costs do not scale linearly at a facility level and Ahanhanzo et al. (2015) demonstrated that the size of a clinic is an important cost driver. In this paper, we examine whether programmatic costs for supplementary immunization activities (SIAs) demonstrate efficiencies of scale. If so, budgeting based on unit cost would drive inappropriate decision making, as shown by Claxton et al. (2016) and suboptimal budget allocation, as previously demonstrated by Fitzpatrick and Bauch (2011).

Previous studies (Walker et al. 2004; Wolfson et al. 2008; Portnoy et al. 2015) have recognized the importance of reporting both fixed and variable costs, as well as differences between subgroups of countries, for example World Bank income class and World Health Organization (WHO) region. However, these studies generally only provide an average per-unit cost without discussion of whether this value is applicable across a range of activity levels, for example moving from pilot program to broad deployment, or when building on an already successful platform. The insufficient understanding of the drivers of cost may be one of the reasons that budgets are often quite far off from actual expenditures (13), and Gandhi and Lydon (2014) demonstrated that vaccination programs may be unable to achieve the levels of population immunity necessary for disease control as a result.

With more flexible (non-linear) cost estimation techniques, we may be able to identify situations where vaccination programs should be able to achieve economies of scale, which could be used to improve the cost-effectiveness of interventions, such as malaria and measles elimination programs. To this end, we have examined historical polio program costs and found that marginal costs vary depending on the scale of the program and that there are efficiencies of scale (costs grow sub-linearly with scale). This supports the move toward using “cost functions” rather than unit costs as the primary method for estimating programmatic budgets.

To examine the question of whether costs grow sub-linearly (demonstrating efficiency of scale), we examined cost data from the Global Polio Eradication Initiative (GPEI). While the GPEI has been unique in its scale to date, other programs such as the Global Fund to Fight AIDS, Tuberculosis, and Malaria (Global Fund) and the Measles and Rubella Initiative (MRI) are both infectious disease-focused programs with international scope. These and other programs targeting disease elimination will likely rely at least in part on SIAs to reach every last child and on strong disease surveillance to measure progress; thus, having a better understanding of how these costs grow will be useful in ensuring programmatic success.

Gandhi and Lydon (2014) reported that operational costs for SIAs administering oral polio vaccine (OPV) range between $0.02-$1.97 per dose in 2010 US dollars, depending on the country. We seek to extend this work by assessing whether average per capita costs for a given country are consistent as the program scales up the number of OPV doses being delivered. The GPEI has also begun legacy planning for post-eradication, capturing lessons learned that can be translated to other diseases. We seek to support that goal by uncovering cost drivers and understanding scale efficiency that are relevant to other diseases being considered for elimination or eradication.

## Data and Methods

The GPEI provides an ideal setting to investigate the linearity of cost data as it includes 12 years of budget data, costs are reported across several operational subcategories, and it includes 86 countries or territories, which comprise the majority of the global expenditures in low- and middle-income countries. With GPEI having organized the polio eradication effort since 1988, we believe there is sufficient internal consistency to facilitate a comparison of costs over time. The data contains budgets for the vaccination and surveillance programs for each country and year, so each country contributes a maximum of 13 data points (one for each year 2005-2017).

This analysis utilizes historical budget data published in the GPEI financial resource requirement (FRR) reports from 2005-2017. Each FRR document contained total GPEI fundraising budgets for high risk countries by cost category. Each FRR also included surveillance and technical assistance costs for all countries involved in the program (including both high and low risk countries), risk classification by country, and SIA schedules. The data was extracted from the published PDFs into a Microsoft Excel workbook. All budgets were reported in millions of United States dollars (USD).

Surge, other, and indirect costs were excluded from the costs totals because they were not consistently reported over time. All other costs were grouped as follows for this analysis.

- OPV – Defined in the FRR as including procurement and transport to the country.
- Vaccination Delivery – A sum of the FRR categories of national or subnational immunization days (NID/SNID) Operations and Social Mobilization.
- Surveillance – The FRR category of acute flaccid paralysis (AFP) Surveillance and Lab, defined as running costs for conducting active surveillance and the laboratory staffing, overhead, and sample processing costs.
- Total – Sum of the three above, plus the FRR category of Technical Assistance.

Budgets from prior to 2017 were adjusted for inflation into 2016-equivalent dollars using the World Bank’s gross domestic product (GDP) deflators (15). The country-specific GDP deflator was used to adjust in-country expenses including operations, social mobilization, technical assistance, and surveillance costs. OPV costs were inflated based on the European Union GDP as a proxy for international developed-world inflation, since these prices are determined in the international pharmaceutical market, not the local economy.

Active surveillance data was obtained from the WHO’s surveillance database, which reports the number of acute flaccid paralysis (AFP) cases by country by year. AFP is the sudden onset of paralysis in children under 15 and is the hallmark symptom of polio. Globally, every AFP case is supposed to be reported and have stool samples collected and tested for poliovirus. This has been the foundation of polio surveillance and is the predominant cost of the GPEI’s surveillance program.

Population totals by age group were sourced from the World Bank (16) and then multiplied by FRR-reported SIA campaign schedules to estimate the number of SIA-targeted individuals.

To explore whether budgeted values are a realistic approximation for actual expenditures, we compared GPEI’s actual expenditures for calendar year 2016 to the budgeted amounts from the 2016 FRR by using a Pearson correlation on both the natural and natural-log scale. Actual expenditures accounting was performed by GPEI and the data was provided to the authors for analysis courtesy of the Bill and Melinda Gates Foundation. Unfortunately, the actual expenditures are not currently available with country-level detail prior to 2016, so that was the only year in which we were able to compare actuals to budgets; however, GPEI has made the total program expenditures are available on their website (17) for years 2013-2016.

We accounted for the repeated observations by country by using a linear mixed effects regression (LMER) model. Specifically, we expect there to be year-on-year correlation in error within countries, since we are dealing with budgetary values that tend to be adjusted relative to the previous year’s values, rather than independently determined every year.

The LMER model has the form log(y_it_) = β_i0_ + β_0_ + β_il_ * log(x_it_) + β_1_ * log(x_it_) where y_it_ is the cost and x_it_ is the number of units for country i in year t and both values have been transformed to the natural-log scale. The β_i0_ and β_i1_ are mean-zero random intercepts and random slopes, respectively and describe the country-specific deviations. The resulting coefficients β_0_ and β_1_ were used to determine the rate at which costs scaled with program size. With this model, a β_1_ that is less than 1.0 means that an increase in x_it_ (e.g., number of vaccine doses distributed) is associated with a decreasing marginal increase in y_it_ (e.g., vaccination delivery cost), implying an efficiency of scale. That is, if the mean-zero random effects are removed, log(y_it_) = β_0_ + β_1_ * log(x_it_) and 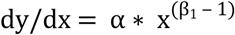 where α = β_1_ * exp(β_0_). Thus, if 0 < β_1_ < 1, then dy/dx decreases as x_it_ increases.

All model fits were done in the software package R (18) version 3.4.2. We used the lmer() function from the lme4 library (Bates et al. 2014) to fit the LMER model.

## Results

We first examined whether a natural-log scale provided a good basis for analysis. Figure 1 displays the natural log transformation of the independent variable (SIA-targeted children, population under age 5, and AFP count) and dependent variable (cost), with each point representing a country-year combination (i.e. countries contribute multiple data points, one for each year it is included in the data set).

**Figure 1:**
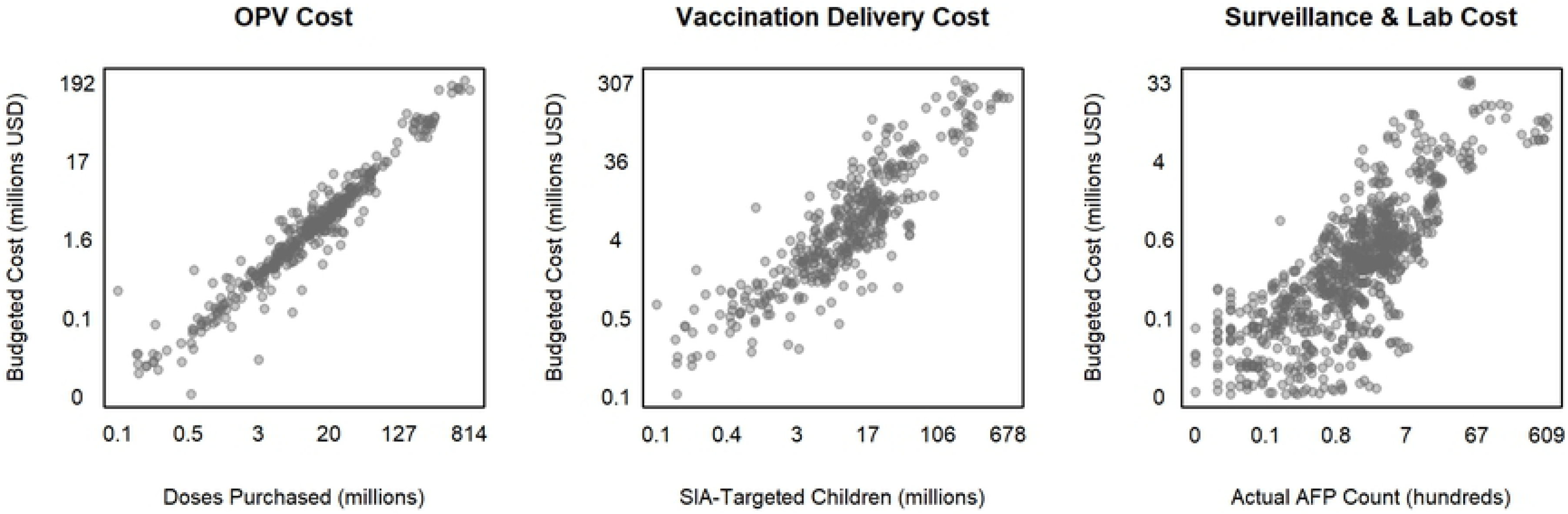
Inflation-Adjusted Costs. Each cost category as a function of its relevant cost drivers (x). Both the independent and dependent variables are plotted on the natural log scale.

To determine whether budgets are a reasonable approximation of actual expenditures, we compared the total 2016 budget and actual expenditures using a Pearson correlation, with the resulting correlation statistic of 0.995. To assess whether the difference in scale across countries was driving the correlation we also calculated the correlation on the natural log scale, with the resulting correlation statistic of 0.935. From this, we concluded that using the historical budgeted data was adequate for our purposes. The remainder of the results in this paper are based on natural log transformed values and utilize only the budget dataset from the GPEI’s FRRs.

Next, we examined the relationship between several hypothesized cost drivers and subsets of costs. The cost categories of “surge” and “other” are not consistently reported over time or for every country, so they were included in the total cost numbers but were not considered independently. The cost categories of “technical assistance” and “indirect” were not considered independently because they are less directly related to the specifics of how a country’s program is being run. Cost values that were considered included:

- OPV procurement cost vs. number of vaccine doses ordered
- Vaccination delivery cost vs. number of SIA-targeted individuals
- Surveillance and laboratory cost vs. actual AFP count
- Total cost vs. population under 5

To assess whether there are efficiencies of scale, we fit the LMER model to each x-y pair to determine the value of β_1._

The LMER model was fit for all four scale-driver pairs. The total cost per capita is provided for informational purposes, since this is how budgets for many programs are currently reported. The OPV vaccine prices are negotiated between UNICEF and vaccine manufacturers on a per-vial basis, this provides a natural control, which should have a β_1_ close to 1.0. Indeed, we find that the confidence interval for β_1_ includes the value 1.0, which provides us with confidence in the model (see Table 1).

**Table 1:** Model Fit Comparisons. LMER country-specific models were fit to historical GPEI budgets from 2005-2016. All values in this table are reported in the units of log(Millions USD). β_1_ column indicates the mean value and their 95^th^ percentile confidence intervals. B_i1_ column indicate the first quartile, median, and third quartile of the country-specific values.

| Cost (Y) | $\beta_1$ log(M USD) | $\beta_{i1}$ log(M USD) |
| --- | --- | --- |
| OPV | 0.97 (0.92, 1.03) | -0.03 / 0.00 / 0.05 |
| Vaccine Delivery | 0.71 (0.61, 0.81) | -0.09 / 0.02 / 0.11 |
| Surveillance | 0.02 (-0.13, 0.16) | -0.23 / 0.03 / 0.27 |
| Total | 0.81 (0.01, 0.91) | -0.10 / -0.02 / 0.11 |

In the remainder of this paper we focus on the two costs over which GPEI has decision making control: vaccination delivery and surveillance. Since a value of less than 1.0 implies an efficiency of scale, the results demonstrate that there are economies of scale for these two costs.

In Figure 2, the lines represent the country-level LMER model fits. Countries are color coded by risk level based on the number of years they were classified in each risk category: endemic (for all 14 years), recurrent (at least three) outbreaks, at least one year of high risk, or low risk.

**Figure 2:**
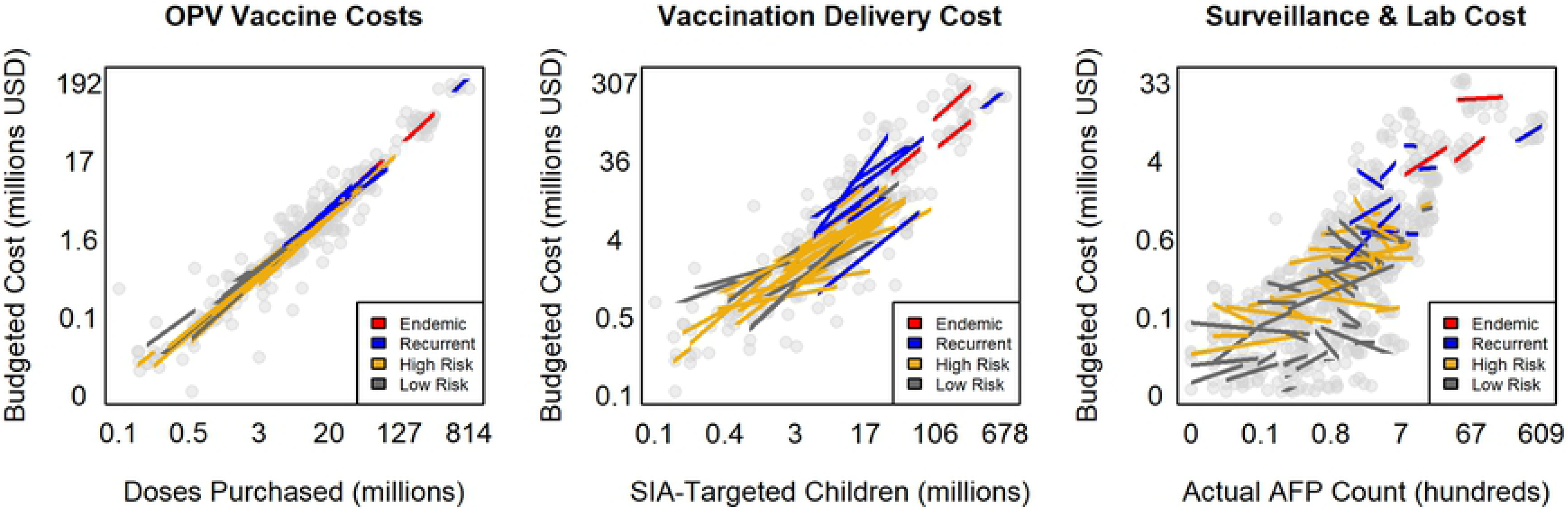
Graphical Model Fits. The three plots show a comparison of the model fits for vaccination delivery costs vs. SIA-targeted individuals and surveillance costs vs. AFP count. Each data point represents a single country-year pair. The LMER fit lines each represent a single country and are drawn to cover the x-and y-range of values found in the actual historical data for that country and are color-coded by risk level.

In Figure 2, we examine the relationship between the number of vaccination doses delivered and vaccination costs and between the number of AFP cases and surveillance costs. The model of surveillance costs varies substantially by country, which suggests that intra-country variation is an important effect. The small magnitude of the slope implies that total surveillance costs are mostly fixed (overhead) and that variable costs are relatively low. The wide range in the country-specific slopes (1^st^ and 3^rd^ quartiles ranging from −0.23 to 0.27) implies that surveillance cost growth is country-specific. The large, mostly endemic and outbreak-prone countries have slopes that demonstrate economies of scale as their surveillance programs have intensified over time. In contrast, the smaller and lower risk countries tend to have flatter (and even sometimes negative) slopes. This implies that surveillance costs vary significantly between countries.

## Discussion

Parameter estimates for vaccination delivery and surveillance costs significantly less than 1.0 support the conclusion that there are efficiencies of scale. These results are consistent with the results of Portnoy et al. (2015), which found that a country’s size (measured as a binary of whether the population exceeded 10 million people) was a statistically important driver of costs, suggesting that smaller countries spend a larger portion of their budget on overhead and achieve less efficiency of scale. It also suggests that published literature should be careful to avoid only reporting the average cost of a health intervention, since it is unlikely that the cost per person is consistent at all scales. It would be preferred if total, fixed, and variable cost categories were reported, as well as known cost drivers (such as number of vaccine doses delivered), to support the development of cost functions rather than single point estimates.

For decision makers, the extent to which there is efficiency at scale should be considered when planning how to scale up or down, and when to intensify disease control efforts. This has implications for how best to roll out an intervention; of course, the epidemiology should drive prioritization but in the context of limited funding, cost considerations can help decision makers to maximize their impact per dollar spent.

Our results are consistent with findings that there are indeed economies of scale in fields as diverse as banking (Din et al. 1996; Wheelock and Wilson, 2017), education (Glass et al., 1995; Abbott and Doucouliagos, 2003), real estate (24), and agriculture (25–27) – although Wilson and Carey (2004) found that optimal scale may be locally-dependent and there is a risk of diseconomies of scale in certain situations (29). There may also be efficiencies of scope by deploying interventions for multiple diseases in an integrated fashion, but we have not assessed that in this paper.

It is interesting to note that while large endemic and outbreak-prone countries have more predictable budgets, this is not always the case for small and low-risk countries, particularly for surveillance costs. The resulting flat- or negatively-sloped model fits do not lend themselves to obvious interpretation and may be indicative that it is more difficult to estimate budgetary needs in low-intensity settings.

While it may be tempting to interpret these results directly, since these models are built on the natural-log scale we cannot draw definitive conclusions about how much cost is fixed vs. variable. This is because the relationship in real terms is of the form 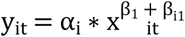 where 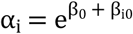. As a result of this structure, the slope α_i_ is dependent on the intercept terms β_0_ and β_i0_, which implies that the rate of change in costs is not consistent across the range of x-values being evaluated. The intercept is also outside the range of our dataset, so direct interpretation of the β_0_ and β_i0_ is not appropriate.

With that in mind, the variation between countries described by their random slopes and intercepts imply that there are important country-specific differences in how costs scale, in addition to the sub-linear scaling up of costs implied by the general model fits. The differences between sub-categories of costs (OPV vs. vaccination delivery vs. surveillance) is also informative and suggest that multiple costs components contribute to scaling efficiencies, which should be further investigated at lower levels of detail than is currently possible from the aggregated budget data that GPEI publishes.

Overall, this analysis suggests a need to move beyond reporting “average per capita” or “average per vaccine” as a standard measure, since they imply a linear growth in costs, without considering whether there may be (dis)efficiencies of scale. Instead, reporting should include the total costs and any known information on the cost drivers, so that other researchers and decision makers can re-use that information more effectively.

By considering the mix of fixed and variable costs, budgeting may become more accurate. Better estimation drives appropriate fundraising and reduces the likelihood that funds are tied up unecessarily, releasing them to be spent on other priorities. This is imperative for elimination programs that are in the process of scaling up, such as regional measles elimination programs and the Central America malaria elimination program.

It is also critical for GPEI to understand these dynamics so that they do not underbudget support for the polio program as it winds down during the certifcation phase after eradication has been achieved. Economies of scale worked in the program’s favor as it grew, with incremental growth requiring less investment than before and overhead costs being spread across a larger base. However, this is also the downside of economies of scale because when a program begins to slow down, the savings may be less than expected (doing half the volume of work does not necessarily reduce the cost by half) and spending levels may remain high even during certification in order to maintain response capacity and a strong active surveillance effort.

While we did find a high correlation between the budget and actual values for the year 2016, the unavailability of prior country-level actual expenditures limits our ability to confirm that this relationship is true historically. However, GPEI does indicate in their 2013-2016 expenditure reports that they consistently spent less than budgeted for reasons including: unanticipated funding from humanitarian or in-country sources, campaign delays or changes in scope, favorable exchange rates, unfilled vacant positions, and vaccine shortage for inactivated polio vaccine (IPV) in 2015-16. Each of these factors contributes to the variance between the budget and actual expenditures, primarily due to changes that could not be anticipated in advance rather than any fundamental misunderstanding of what it costs to implement the annual polio program. With that in mind, the authors believe it is reasonable to use budgeted values from the FRRs as a proxy for true costs for the purposes of this analysis. The analysis could be validated further if actual expenditure data was available in the future to perform the same analysis.

Additionally, the GPEI’s FRR documents are focused on the amount of external funding needed to run the program, so they do not include in-kind costs provided by host countries nor self-funded programs. To address this, we opted to exclude any country-year of data where there was a known bias, but this may result in some selection bias and there may be some remaining unknown variance between GPEI’s budget and total costs.

This model could be further expanded by including explanatory variables such as development indicators, health system infrastructure, and disease program goals (i.e. endemic, disease control, elimination, eradication) in order to enhance the predictive power of the model and assess which (if any) are drivers of cost. The same type of exploration that has been done in this paper would also be useful for non-immunization programs such as mass drug adminstration, bed net distribution, and other broad-reaching health interventions.

Reporting a more detailed cost framework, combined with a better understanding and research into where there are economies of scale, should help inform funders and operational decision makers about the best way to ensure that adequate resources are mobilized and deployed, to maximize the impact of public health interventions. These lessons about cost efficiencies should be incorporated into future efforts to design cost functions that describe the non-linear way that costs accrue in reality, so that we can translate the experience of polio eradication to other global disease programs.

## Acknowledgements

We thank those who collaborated with us on the development of this manuscript including Xiaodong Cai, Ticky Esoh, Suchita Guntakatta, Hil Lyons, Ann Ottosen, Michiyo Shima, Britta Tsang, Arie Voorman, and Dan Walters.

## Funding

This work was supported by Bill and Melinda Gates through the Global Good Fund. The funders had no role in study design, data collection, data analysis, the decision to publish, or preparation of the manuscript.

## Conflicts of Interest

The authors declare no conflicts of interest.

